# Pattern of Self-Administered Vapor Fentanyl Exposure Determines Long-term Behavior Consequences, in Mice with or without Neuropathic Pain

**DOI:** 10.1101/2022.09.22.508919

**Authors:** Samantha Cermak, Maria Virginia Centeno, Rami Jabakhanji, Andrew Vigotsky, Julia Cox, Andrew Brink, Apkar Vania Apkarian

## Abstract

We studied the behavioral consequences of fentanyl vapor self-administration (SA) in mice with and without chronic neuropathic pain (one month after spared-nerve injury(SNI) model or sham injury). We assessed fentanyl consumption, motivation, and seeking during as well as anxiety, hyperactivity, immobility, and pain for two regimens of fentanyl SA: 1) Dose escalation, where over a 3-week period mice are exposed (daily 2-hour sessions) to escalating numbers of fentanyl puffs per active nosepoke (from 1 puff/active nosepoke for first 3 days, up to 6 puffs/active nosepoke in days 16-18). 2) Effort escalation, where over a 3-week period (daily 2-hour sessions) mice need to increase effort to acquire the same amount of fentanyl (fixed ratio 1 (FR) = 1 active nosepoke results in 1 fentanyl puff, while second and third week we use FR5 and FR10). We observe sex-, injury- and regimen-dependent differences in outcomes. Importantly the dose escalation regimen resulted in higher seeking behavior (post forced abstinence, context and cue driven nosepoking in the absence of fentanyl delivery), long lasting increased anxiety, immobility, and hyperactivity, as well as transient but full pain relief in SNI mice. Therefore, this regimen seems a better rodent model for translating outcomes to human chronic pain patients managed with opioids.

## Introduction

Approximately 20% of individuals will experience chronic pain [1-3], propagating to the societal level to create an economic burden of $560–635 billion [4]. Notwithstanding its massive societal impact, limited effective therapies exist for chronic pain, and 20% of chronic pain patients report prescription opioid use [5]. However, chronic opioid use has several side effects, including the risk of addiction [6]. This is especially true for long-term opioid use in chronic pain [7]. Approximately 21–29% of chronic pain patients who are on opioids misuse them, and rates of addiction are estimated to be between 8–12% [8]. Thus clinical management of chronic pain is itself a contributor to the ongoing “opioid epidemic”, characterized by overdoses and opioid-related deaths across the country, which are on the rise [9, 10], particularly with synthetic opioids such as fentanyl [9]. To properly address the clinical issues regarding opioid exposure in chronic pain it is necessary to garner better understanding of underlying mechanisms. The latter is contingent on adequate animal models of addiction in combination with chronic pain.

Rodent models for opioid addiction, especially in mice, has been hampered by various technical issues especially regarding self-administration models which traditionally use intravenous catheters [11, 12]. Recent vapor delivery methods have greater promise [13] as they are noninvasive and thus can be readily combined with other manipulations for example with genetically engineered mice. Here we examine the behavioral consequences of exposing mice to multiple weeks of fentanyl using noninvasive vapor self-administration following the technology introduced by Moussawi et al. [13], and compare long-term behaviors for various regimens of fentanyl exposure regarding their correspondence to the humans with chronic pain and using opioids for managing the pain.

The standard clinical opioid management of chronic pain patients begins with dose escalation, to attempt to find the appropriate does for adequate pain relief. Also, when patients develop a tolerance to their opioids, physicians further increase their patients’ dosages to sustain adequate pain relief. Yet, dose escalation does not necessarily provide pain relief [14, 15] and is associated with negative outcomes, such as an increased risk of overdose or opioid-related accidents [16, 17]. Nevertheless, opioid dose escalation is common with long-term treatment, across different types of opioids and routes of administration [16, 18-23].

Despite its ubiquity, few preclinical models of opioid addiction have addressed dose escalation [24, 25]. More commonly, opioid studies in rodents have studied effort escalation, a method used to assess extent of motivated drug consumption in rewarding drugs such as cocaine. In this method, animals are trained on a fixed-ratio (FR) schedule, usually at FR1, so one action leads the animal to receive a given amount of drug [26, 27]. Sometimes this entails an increasing amount of “effort”, either nosepokes or lever presses, to receive the same amount of drug [13] to test the amount of effort an animal will spend to acquire the drug.

Opioid self-administration studies using a FR schedule have shown that arthritic rats, compared to sham, self-administer fentanyl at greater levels, mediated not by reward or dependence but by the reversal of hypersensitivity [28, 29]. Similarly, for animals with spinal nerve ligation, a model of neuropathic pain, intravenous opioid self-administration was reduced for low doses, but high opioid doses in these animals were maintained, and correlated with the alleviation of hypersensitivity [12]. However, while using vapor fentanyl self-administration has been established [13], how chronic neuropathic (SNI) animals behave has not been explored, especially for different regimens of opioid exposure.

Since dose escalation more closely resembles the clinical tradition of pain management in chronic pain, we studied its behavioral consequences in comparison to effort escalation, in chronic neuropathic mice (spared-nerve injury model) or sham. Opioid seeking and behavior were compared between animals undergoing a dose escalation regimen and a typical effort escalation regimen. We found that animals undergoing dose escalation show persistent seeking behavior and behavioral differences, especially compared to effort escalation. Particularly, SNI animals undergoing dose escalation exhibited long-term (in the absence of fentanyl) lower mobility, increased anxiety, increased inactivity, and transient analgesia. These properties better match the human condition of chronic pain managed with opioids. Thus, vapor fentanyl self-administration dose escalation seems to be a better preclinical model for studying opioid use dependence especially in chronic pain.

## Methods

### Subjects

A group of male (n=8) and female (n=8) adult C57BL/6 mice (mass =17–30 g) were used for the first cohort of dose and effort escalation experiments. These animals first underwent the effort escalation regimen, and following three weeks of rest, underwent the dose escalation regimen. A second cohort of eight naïve, healthy mice, both male (n=3) and female (n=5) was used exclusively for the dose escalation fentanyl self-administration analysis. Animals were group housed and given free access to both food and water. Animals were housed on a 12/12-hour light/dark cycle at 72 ± 2°F temperature and 50–70% humidity. All testing was conducted during the dark cycle. Animals were handled regularly to reduce stress. All studies adhered to ethical guidelines and were approved by the Animal Care and Use Committee of Northwestern University.

### Spared Nerve Injury (SNI)

Subjects of the first cohort received surgery for either a spared nerve injury (SNI) (n=8) or sham (n=8) [30]. Animals were anesthetized with isoflurane (1.5–2.5%) in a mixture of 30% N_2_O and 70% O_2._ In animals receiving a SNI, the sciatic nerve was exposed at the trifurcation of peroneal, tibial, and sural branches on the left leg. The common peroneal and tibial nerves were cut and tied with 6-0 vicryl sutures, while the sural nerve was left intact. In animals receiving sham surgery, the sciatic nerve was simply exposed and left intact.

### Fentanyl Self-Administration Apparatus

A Plexiglass chamber (14cm × 20cm × 23cm) was used for fentanyl vapor self-administration, modeled after Moussawi et al.[13]. Chambers were placed inside a sound-attenuating cabinet (Med Associates) to minimize light and noise, and the cabinets were ventilated with fans. The chamber was equipped with nosepokes on either side of the chamber. A negative pressure system sucked the vapor into the chamber when the vaporizer was active. The vapor flow was regulated so the vapor was cleared within 60 seconds. The rewards, active and inactive nosepokes, and their timestamps were recorded by a computer using Med Associates software.

### Drug

5 mg/ml fentanyl HCl was dissolved in a vehicle consisting of 20% propylene glycol and 80% glycerol. Fentanyl was obtained from National Institute on Drug Abuse (NIDA), Intramural Research Program Pharmacy, Baltimore, MD, USA.

### Effort Escalation

#### Program

During the 2-hour self-administration period, all lights were initially off. A nosepoke to the active port would result in vapor fentanyl delivery in the chamber for two seconds, and the active nosepoke port light would turn on for one minute. Following a vapor delivery, there was a one minute time out period, in which any further active nosepokes would not result in drug delivery. After this period, the active nosepoke light would turn off. The inactive nosepoke light remained off, and any pokes to the inactive port were inconsequential.

#### Acquisition

Animals were trained for eight days on an FR1 schedule, in which one nosepoke resulted in one puff of fentanyl. Following this period, animals were given a break of seven days and then reintroduced to FR1 for three days. After two days of rest, they completed the same self-administration task on an FR5 schedule for five days. Then, after another two days of rest, they self-administered on an FR10 schedule for five more days, which concluded the acquisition phase.

#### Cue-Induced Seeking(CS)

After 2, 7, and 14 days of abstinence, we administered a 2-hour drug-seeking test. During the drug-seeking task, no fentanyl puffs were given, although the same drug cues were present as acquisition.

### Dose Escalation

#### Program

During the 2-hour self-administration period, both lights were on at the start of the session. A nosepoke to the active port would result in vapor fentanyl delivery in the chamber for 2 seconds, and the active nosepoke port would blink on and off while the inactive light would turn off. During this time, there was a one-minute time-out period in which further active nosepokes would not result in drug delivery. Inactive port nosepokes were inconsequential.

#### Acquisition

Animals underwent the dose escalation regimen schedule for 18 consecutive days, during which time there was 1 session per day and 2 hours per session. During the dose escalation regimen, testing was on an FR1 schedule, but with increasing the amount of fentanyl delivery for every successful nosepoke. Starting with 1 puff (1P) per reward, we increased the number of puffs per reward by one every three days, i.e., 1P, 2P, 3P, 4P, 5P, and 6P. There was no maximum drug delivery.

#### ue-Induced Seeking(CS)

A 2-hour drug-seeking test was given following seven days(n=16) or 14 days(n=8) of abstinence. Again, no fentanyl was administered during the seeking task, but the same cues were present.

### Open Field

For open field testing [31, 32], we placed a Arducam for Raspberry pi 1080p HD monochrome camera above each of the wide Plexiglass arena (40cm × 40cm × 40cm). We recorded video at a rate of 30 frames per second. The six were connected to a computer and recorded using Streamlabs, an open-source video recording software. The mouse was free to explore the open field arena during the five-minute testing period. The videos obtained were analyzed in a fully automated process using ANY-maze Video Tracking System software (Stoelting Co.), Python, and R software. Parameters of interest included distance (meters), the time in the periphery (percentage), and the amount of time spent inactive (seconds). Measures were collected five minutes following testing (herein “5m”) and one day following (herein “1d”). The 1d testing was a surrogate for a baseline without acute fentanyl exposure, considering any effects of prolonged fentanyl use. Open field was collected at baseline before any testing, during each stage of self-administration, and during abstinence. In dose escalation, these stages included baseline (BL), 1P, 2P, 3P, 4P, 5P, 6P, and CS. For effort escalation, we collected open field during BL, FR1, FR5, FR10, and CS. Open field testing was recorded for five minutes, except for one continuous 2-hour open field conducted to assess the time-course of fentanyl.

### Mechanical Sensitivity

Animals were placed in a Plexiglass box with a grid floor and habituated to the box for 45 minutes. The mechanical allodynia thresholds were measured using MouseMet eVF electronic von Frey (Topcat Metrology Ltd, UK). The apparatus measures the force (grams) at which the animal withdraws the paw. Each rear paw was assessed three times and then averaged[33]. The single threshold obtained, expressed in grams, provides a quantitative assessment of the mechanical allodynia response. Animals were tested before each treatment to obtain a baseline, and then the animals were reassessed after treatment at various stages. In dose escalation, these stages included BL, 4P, and abstinence. For effort escalation, these stages included BL, FR1, FR5, FR10, and abstinence. von Frey recordings were generally collected approximately 45 minutes to 1 hour following fentanyl self-administration, except for one continuous von Frey. During this continuous von Frey session, testing was conducted starting immediately after fentanyl self-administration, consisting of eight trials approximately ten minutes apart.

### Data Analysis

We used generalized linear mixed-effects models with Poisson and negative binomial (in the case of overdispersion) likelihoods and log link functions to evaluate the effects of stage and injury on nosepoke and reward behavior [34]. Four models were created for each regimen (i.e., effort and dose escalation) with the following dependent variables: (1) the number of active nosepokes during each session of the acquisition phases; (2) the number of puffs (rewards) during each session of the acquisition phases; (3) the total number of puffs (rewards) over the entire acquisition phase; and (4) the number of nosepokes for each port during seeking. For all models, we parameterized the effects of injury (SNI vs. sham) and sex (male vs. female), with session included as a continuous covariate, and observations were nested within each mouse (random effects). For model (4), we included an interaction term with port (active vs. inactive) to draw inferences concerning the preference for the active port relative to the inactive port.

Repeated measures analyses of variance (RM-ANOVA) were used to compare the effects of stage and injury (SNI vs. sham) for each open field parameter (distance, time in the periphery, and inactivity). Four separate RM-ANOVAs were performed for each outcome, both at BL and cue-induced seeking (CS): (1) effort escalation, 5m after fentanyl self-administration; (2) effort escalation, 1d; (3) dose escalation, 5m; and (4) dose escalation, 1d. Each RM-ANOVA included stage, group, and their interaction as independent variables. For dose escalation, the stages included consisted of BL, 1P, 2P, 3P, 4P, 5P, and 6P, while in effort escalation, the stages consisted of BL, FR1, FR5, and FR10.

RM-ANOVAs were also conducted to assess the effects of stage and injury on von Frey withdrawal thresholds. This was done for each paw (left and right). For dose escalation, von Frey measurements were collected at BL, 4P, and during abstinence. For effort escalation, von Frey measurements were collected at BL, FR1, FR5, FR10, and abstinence.

For the continuous von Frey, we fit a RM-ANOVA with injury and trial(1:8), to assess withdrawal threshold for each paw. For the continuous open field measurements, a repeated measures ANOVA was used with injury and trial (1:12) on each distance and percent time in the periphery.

## Results

### Effort Escalation (figure 1a-f)

During the acquisition phase of effort escalation, active nosepokes increased with increasing FRs (*P* < 0.001), but female SNI mice increased active nosepokes (exhibited higher motivation) more than the other three groups (male and sham mice) (*P* < 0.001) (Figure 1a). As a result, female SNI mice were able to maintain similar fentanyl doses across acquisition stages (FRs), whereas other mice decreased fentanyl consumption with increased FR (*P* < 0.001) (Figure 1b). Despite this, all consumed similar amounts of fentanyl over the course of acquisition (all *P* > 0.05) (figure 1c). When mice were reintroduced to the fentanyl boxes after withdrawal (i.e., for the cue/context driven seeking task in the absence of fentanyl), total nosepokes (figure 1d) and active port preference (active nosepokes – inactive nosepokes, figure 1e) were higher during seeking in comparison to during acquisition (*P <* 0.001), and female mice and SNI mice had a greater preference for the active port, suggesting higher vulnerability for opioid dependence (*P <* 0.001), (*P* < 0.001 for interaction) (figure 1f).

**Figure 1:**
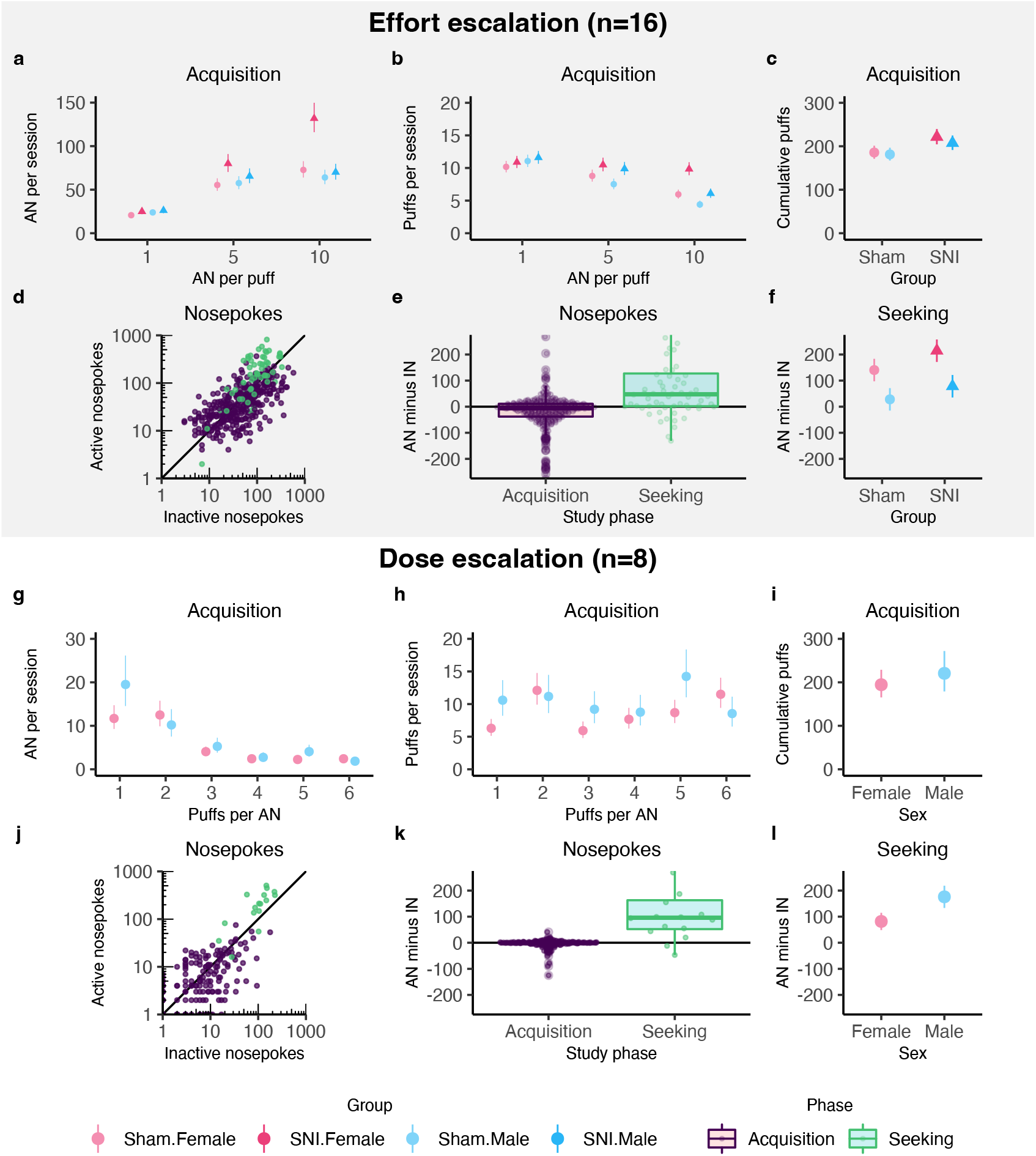
Comparison of fentanyl vapor self-administration in dose escalation and effort escalation. (a-f) Effort escalation vapor fentanyl self-administration. (a) Mean ± SE active nosepokes averaged across all sessions. (b) Mean ± SE fentanyl puffs received averaged across all sessions. (c) Mean ± SE cumulative fentanyl puffs received over the course of the experiment. (d) Active and inactive nosepokes by animal during acquisition and seeking in abstinence. (e) Preference of active port to inactive port by animal. (f) Mean ± SE relative preference during cue-induced seeking task. (g-l) Dose escalation vapor fentanyl self-administration. (g) Mean ± SE active nosepokes averaged across sessions. (h) Mean ± SE fentanyl puffs averaged across sessions. (i) Mean ± SE cumulative fentanyl puffs received over the course of the experiment. (j) Active and inactive nosepokes by animal during acquisition and seeking in abstinence. (k) Preference of active port to inactive port by animal. (l) Mean ± SE relative preference during cue-induced seeking task.

### Dose Escalation (figure 1g-l)

During the acquisition phase of dose escalation, the mice had fewer active nosepokes with increasing doses (*P* < 0.001) (figure 1g). However, the decrease in active nosepokes was approximately inversely proportional to the increase in dose, resulting in similar fentanyl consumption across the acquisition phase (Figure 1h). Mice consumed similar amounts of fentanyl over the course of acquisition (all *P* > 0.05) (figure 1i). When mice were reintroduced to the fentanyl boxes after withdrawal (i.e., for the seeking task), total nosepokes (figure 1j) and active port preference (active nosepokes – inactive nosepokes, figure 1k) were much higher during seeking in comparison to during acquisition (*P <* 0.001), and male mice had a greater preference for the active port (*P <* 0.001) (figure 1l).

### Open Field: Mobility (figure 2)

There was an effect of fentanyl acquisition stage (e.g. 1P,2P…6P or FR1, FR5, FR10) on distance traveled as measured 5 minutes following fentanyl SA in both dose escalation (*F*(6,97) = 26.910, *P* < 2×10^−16^) (figure 2a) and effort escalation (*F*(3,55) = 24.152, *P* = 4.2×10^−10^) (figure 2b). The overall increased hyperactivity immediately after opioid exposure in both regimens implies development of opioid dependence. With dose escalation hyperactivity continues to increase, while with effort escalation hyperactivity decreases. Although we see a similar trend of hyperactivity being higher in sham vs SNI during opioid acquisition 5 minutes after SA, there was only a significant interaction of injury (e.g. SNI or sham) and stage on distance traveled in the dose escalation (*F*(1,97) = 2.198, *P* = 0.0496) (figure 2a), but not effort escalation group (*F*(1,55) = 1.379, *P* = 0.026) (figure 2b). There was an effect of stage (*F*(6,97) = 9.065, *P* = 6.91×10^−8^) and injury (*F*(1,97) = 19.548, *P* = 2.56×10^−5^) on distance traveled 1day after fentanyl SA in the dose escalation group (Figure 2a), but not in the effort escalation group (*F*(1,55) = 0.161, *P* = 0.690; *F*(3,55) = 2.141, *P* = 0.106) (figure 2b). We consider behavior 1 day after fentanyl SA a state where the animals are devoid of opioids, thus these behaviors are long-term aftereffects of opioid exposure. Thus, long-term mobility decreases with dose escalation but remains unchanged with effort escalation. There was a significant interaction of group and stage on distance traveled between baseline and cue-induced seeking in dose escalation (*F*(1,27) = 6.424, *P* = 0.0174) (figure 2c), but not effort (*F*(1,27) = 1.986, *P* = 0.170) (figure 2d). The latter is also a long-term aftereffect of fentanyl exposure.

**Figure 2:**
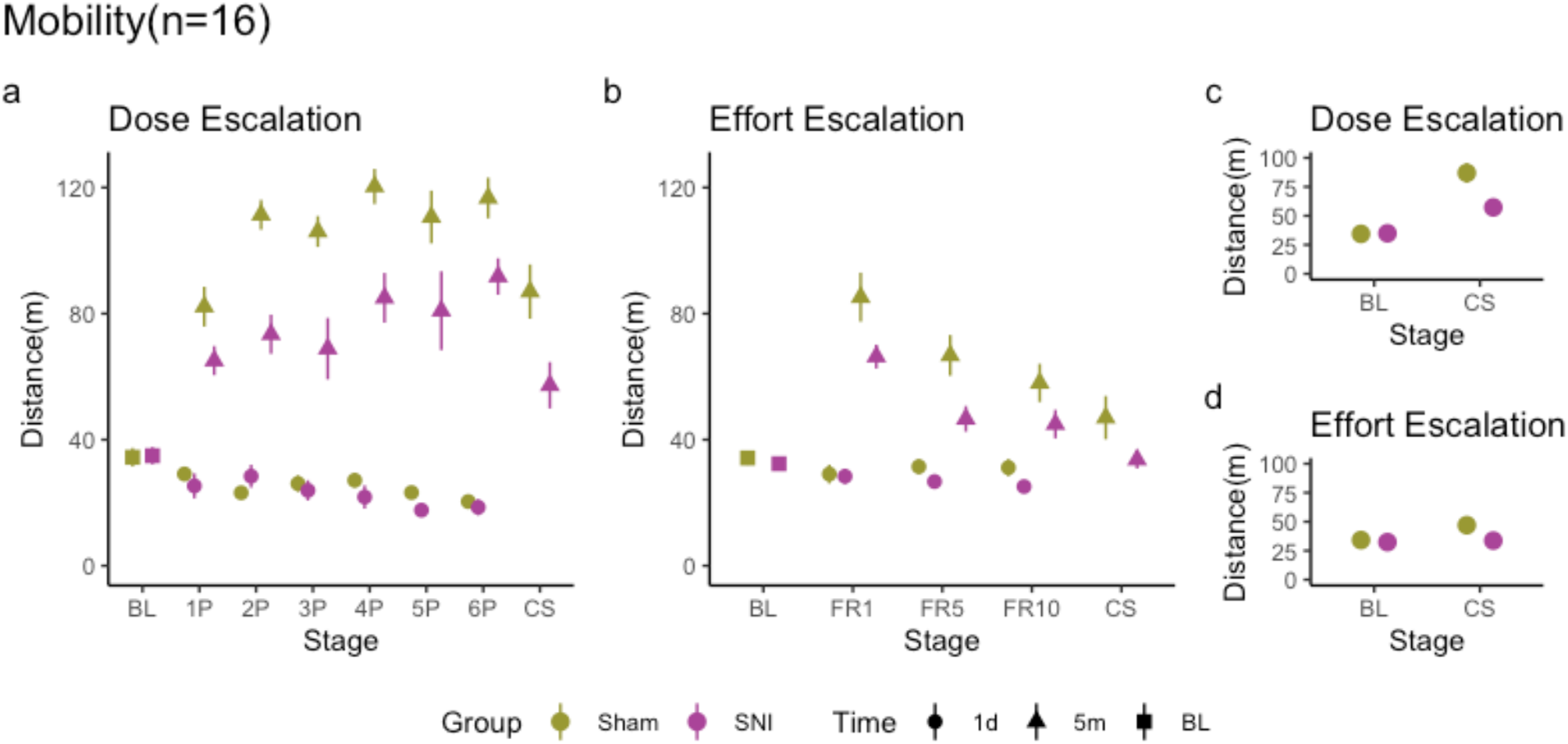
Fentanyl use increases mobility in animals and differentially increases mobility in animals with SNI. (a) Dose escalation mobility data. Mean ± SE distance traveled either 5 minutes (5m) or 1 day (1d) following fentanyl self-administration. Stage indicates time of testing: baseline (BL), fentanyl (1P:6P,P=puffs per active nosepoke), or following a cue-induced seeking test in abstinence (CS). (b) Effort escalation mobility data. Mean ± SE distance traveled either 5 minutes (5m) or 1 day (1d) following fentanyl self-administration. Stage indicates time of testing: baseline (BL), fentanyl (FR1, FR5, FR10), or following a cue-induced seeking test in abstinence (CS). (c) Dose escalation mobility at baseline compared to cue-induced seeking test. (d) Effort escalation mobility at baseline compared to cue-induced seeking test.

### Open Field: Anxiety (figure 3)

There was an effect of stage on percent time spent in periphery of open field (a surrogate measure of increased anxiety) 5 minutes after fentanyl SA in the dose escalation group (*F*(6,97) = 32.687, *P* < 2×10^−16^) (figure 3a) and effort escalation (*F*(3,55) = 7.023, *P* < 0.001) (figure 3b). However, with dose escalation anxiety measure continues to increase, while with effort escalation it increases at FR1 and then reverts towards baseline levels at higher FRs. There was only an effect of stage on percent time in periphery 1 day (long-term aftereffect) post-fentanyl SA in dose escalation (*F*(6,97) = 2.859, *P* = 0.013) (figure 3a), not effort escalation (*F*(3,55) = 1.207, *P* = 0.316) (figure 3b). For most doses, this 1 day post-fentanyl increased anxiety was more pronounced in SNI mice. There was an effect of stage on the percent of time spent in the periphery between baseline and cue-induced seeking in dose escalation (*F*(1,27)=7.379, *P* = 0.011) (figure 3c), but not effort escalation (*F*(1,27)=0.502, *P* = 0.485) (figure 3d), although there was a significant interaction of injury and stage (*F*(1,27)= 12.042, *P* = 0.002). Overall, for dose escalation we observe enhanced anxiety immediately after fentanyl SA, one day after, and also during seeking (no fentanyl), while for effort escalation we only observe transient increases in anxiety.

**Figure 3:**
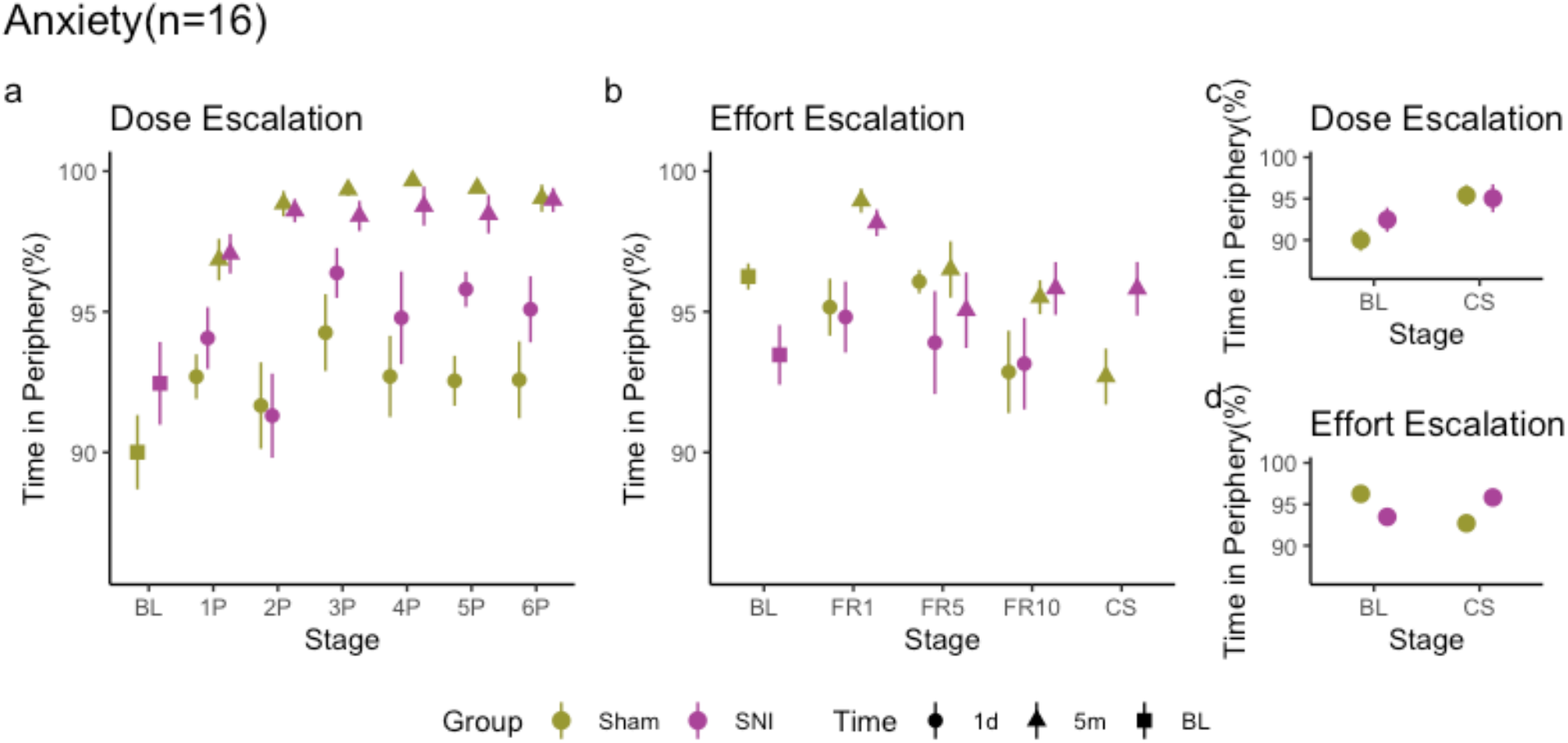
Fentanyl use increases anxiety in dose, but not effort escalation. (a) Dose escalation anxiety data. Mean ± SE distance traveled either 5 minutes (5m) or 1 day (1d) following fentanyl self-administration. Stage indicates time of testing: baseline (BL), fentanyl(1P:6P,P=puffs per active nosepoke), or following a cue-induced seeking test in abstinence (CS). (b) Effort escalation anxiety data. Mean ± SE distance traveled either 5 minutes (5m) or 1 day (1d) following fentanyl self-administration. Stage indicates time of testing: baseline (BL), fentanyl (FR1, FR5, FR10), or following a cue-induced seeking test in abstinence (CS). (c) Dose escalation anxiety at baseline compared to cue-induced seeking test. (d) Effort escalation anxiety at baseline compared to cue-induced seeking test.

### Open Field: Inactive Time (figure 4)

There was an effect of stage on inactive time (immobility) 5 minutes after fentanyl SA in dose (*F*(6,97) = 2.515, *P =* 0.026) (figure 4a), but not effort escalation (*F*(3,55) = 2.197, *P* = 0.099) (figure 4b). However, there was only an effect of injury on inactive time in dose escalation (*F*(6,97) = 12.366, *P* < 0.001) (figure 4a), not effort (*F*(3,55) = 1.192, *P* = 0.280) (figure 4b). There was no effect of stage on inactive time 1d after fentanyl SA dose escalation(*F*(6,97) = 2.133, *P* = 0.056) (figure 4a), but there was on effort escalation (*F*(3,55) = 3.012, *P* = 0.038) (figure 4b). The effects of time from baseline to cue-induced seeking on inactive time was significant for dose (*F*(1,27)=4.226, *P* = 0.0496) (figure 4c), but not effort escalation (*F*(1,27) = 2.939, *P* = 0.098) (figure 4d).

**Figure 4:**
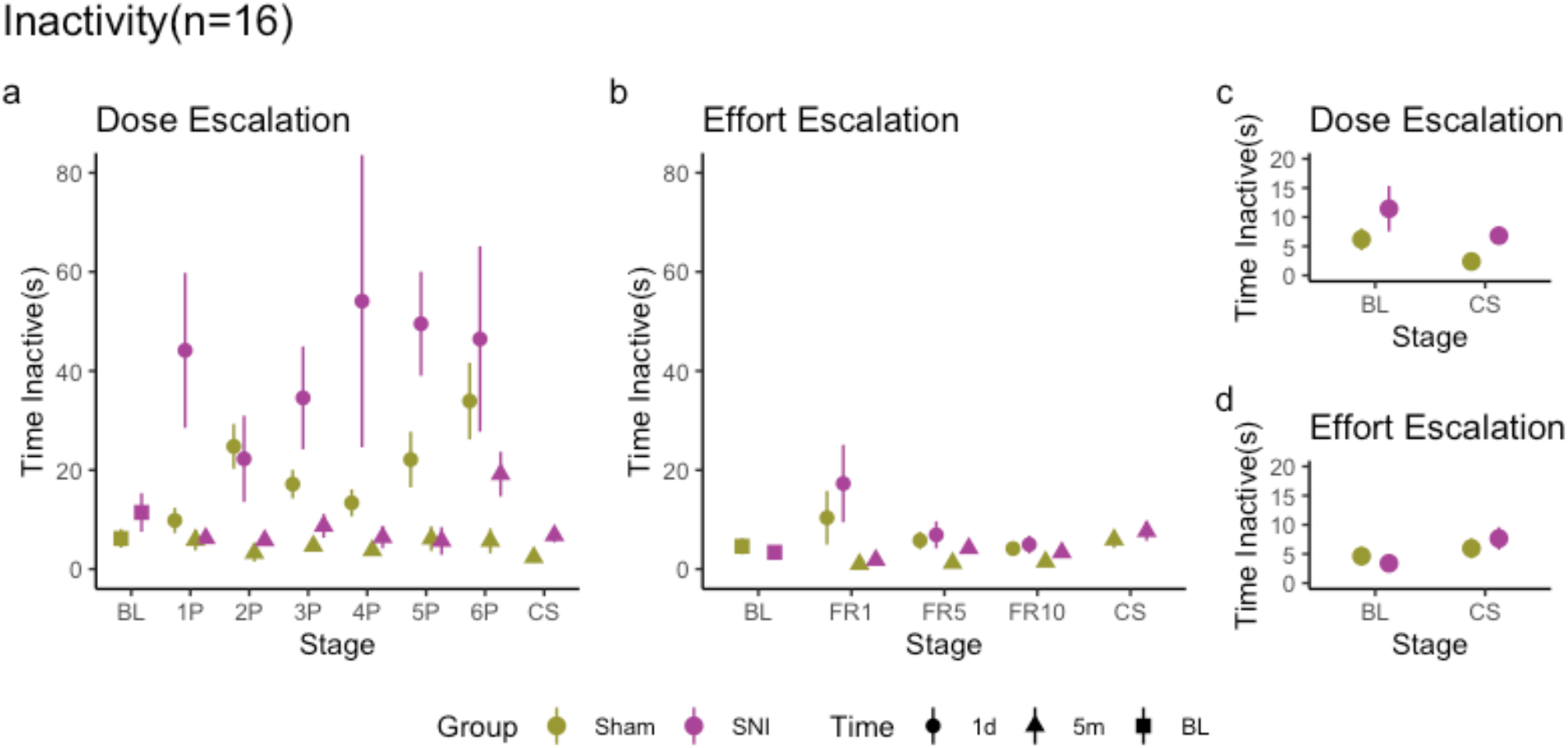
Dose escalation regimen shows increased inactivity 1d, but not 5m after fentanyl use. particularly in animals with SNI. (a) Dose escalation inactivity data. Mean ± SE distance traveled either 5 minutes(5m) or 1 day(1d) following self-administration. Stage indicates time of testing: baseline(BL), fentanyl(1P:6P,P=puffs per active nosepoke), or following a cue-induced seeking test in abstinence (CS). (b) Effort escalation inactivity data. Mean ± SE distance traveled either 5 minutes(5m) or 1 day(1d) following self-administration. Stage indicates time of testing: baseline(BL), fentanyl(FR1, FR5, FR10), or following a cue-induced seeking test in abstinence (CS). (c) Dose escalation inactivity at baseline compared to cue-induced seeking test. (d) Effort escalation inactivity at baseline compared to cue-induced seeking test.

### Mechanical Sensitivity (figure 5)

Mechanical sensitivity or tactile allodynia as a surrogate measure of pain-like behavior was usually measured at 2 hours after fentanyl SA, presumed to be at a time when fentanyl is washed out of blood, thus testing for analgesia (renormalization of tactile allodynia towards baseline levels), or for hyperalgesia (increased tactile sensitivity). There was a significant interaction of stage and injury on withdrawal thresholds for the injured paw for dose escalation (*F*(2,28) = 5.283, *P* = 0.011) (figure 5a), but not for effort escalation (*F*(4,57) = 0.812, *P =* 0.523) (figure 5b). For dose escalation, at a dose of 4p we observe complete analgesia for the injured paw which returns back to allodynia during abstinence (CS, figure 5a, injured paw). There was an effect of stage on von Frey withdrawal threshold in the injured paw for dose (*F*(2,28) = 12.217, *P* < 0.001) (Figure 5a) and effort escalation F(4,56) = 4.665, *P* = 0.003) (figure 5b). However, while there was no significant effects of stage or injury condition on withdrawal threshold in the uninjured paw for dose escalation (figure 5a), there was a significant effect of stage F(4,56)=12.430, *P* = 2.66×10^−7^) and injury x stage in effort escalation (*F*(4,56) = 0.092, *P* = 0.913) (figure 5b). Thus, we observe transient analgesia with dose escalation and sustained hyperalgesia with effort escalation.

**Figure 5:**
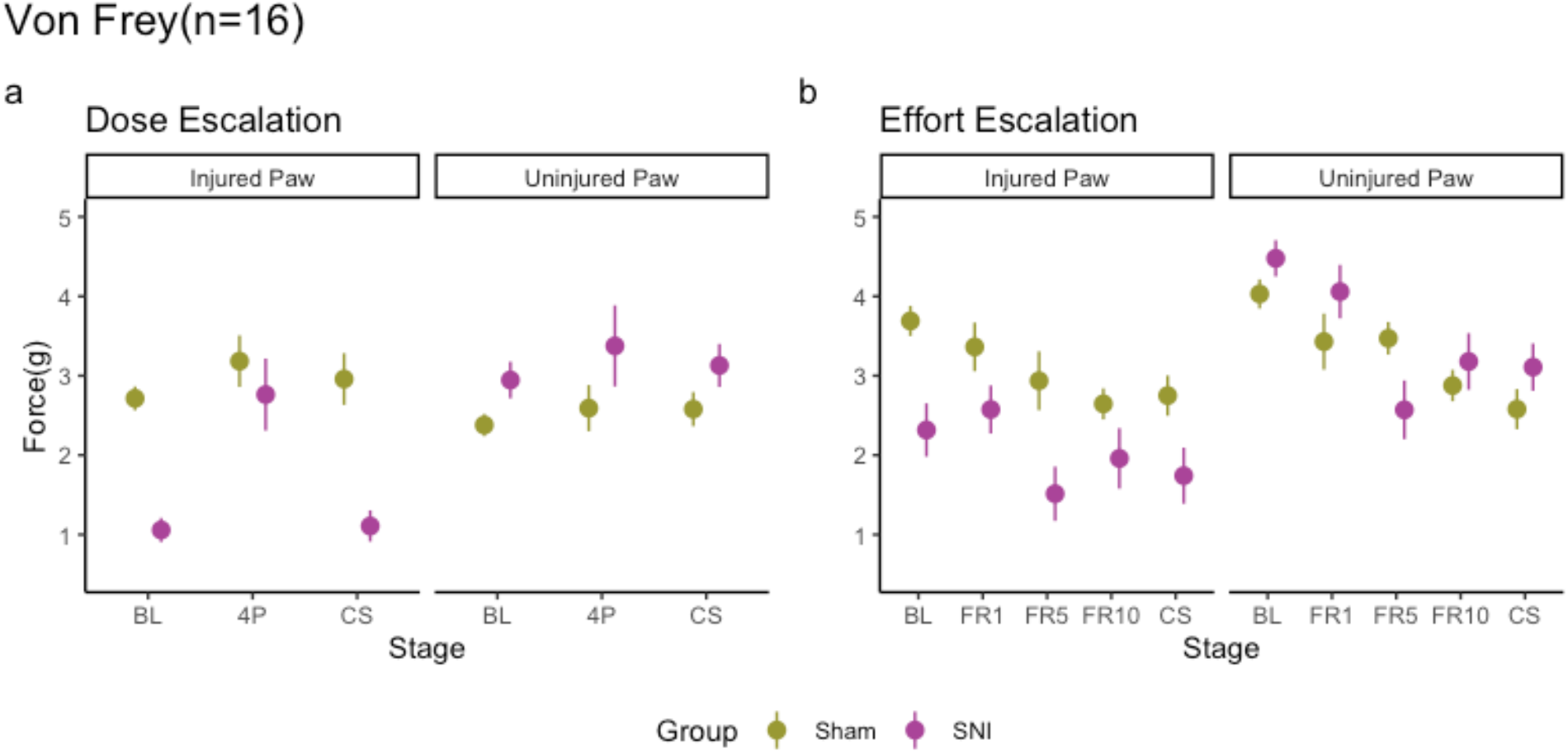
SNI animals in dose, but not effort show hypoalgesia following self-administration, while SNI and sham animals in effort escalation show hyperalgesia. (a)Dose escalation animals with SNI show increased withdrawal threshold to von Frey in the injured but not uninjured paw following self-administration(4P), compared to baseline. Withdrawal threshold returns to the same levels of baseline during following a cue-induced seeking test in abstinence (CS). (b)Effort escalation animals with both SNI and sham show decreased withdrawal threshold following self-administration(FR1, FR5, FR10) and 7d after following a cue-induced seeking test in abstinence (CS).

### Continuous Open Field and Mechanical Sensitivity (figure 6)

In the dose escalation mice, we tested time dependent changes in mobility (hyperactivity), time in periphery (anxiety), and tactile allodynia (pain) starting at 5 minutes after fentanyl SA and over the next 2 hours (figure 6). There was an effect of trial on distance traveled (*F*(11,167) = 30.255, *P* < 2×10^−16^) (figure 6a). There was an effect of injury (*F*(11,167) = 4.597, *P* = 0.034) and trial (*F*(11,167) = 2.094, *P* = 0.023) on percent time in periphery (Figure 6b). Both hyperactivity and anxiety decreased in time, but anxiety in SNI decreased more slowly. Over the course of 8 trials, there was a significant effect of injury on the withdrawal threshold for the injured paw (*F*(1,14) = 7.779, *P* = 0.145), but not the uninjured paw (*F*(1,14) = 0.596, *P* = 0.453) (figure 6c). For the injury side, we observe both sham and SNI exhibit initial hypoalgesia (pain relief above baseline), and in both groups by trial 8 pain levels match those for baseline in sham, ie the SNI animals continue to exhibit analgesia compared to their baseline level of allodynia.

**Figure 6:**
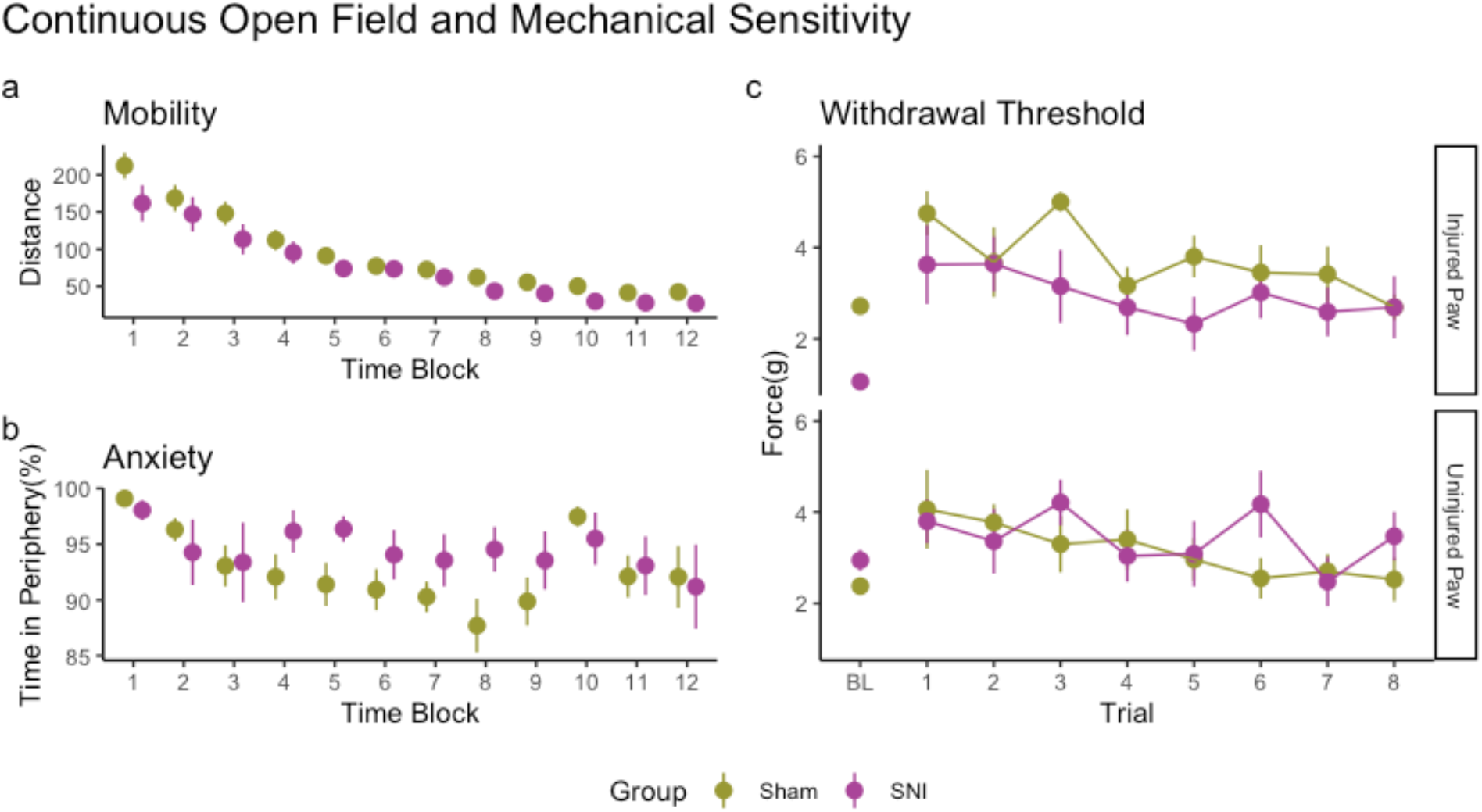
The effects of fentanyl on mobility, anxiety, and withdrawal threshold gradually return to baseline over a 2-hour period. (a) Mean ± SE distance traveled over a continuous 2-hour period, 12 timeblocks of 10 minutes each. All animals show increased mobility early on. (b) Mean ± SE time in periphery shows SNI animals have increased lasting anxiety compared to sham animals.(c) von Frey threshold hypoalgesia is higher in earlier trials and lowers over time.

## Discussion

These results indicate that the pattern of fentanyl self-administration exposure influences behavior, namely, drug seeking, open field, and mechanical sensitivity. Although both regimens included the same concentrations of fentanyl vapor and animals consumed similar quantities of total fentanyl, behavioral differences emerged. Dose escalation produced lasting effects, including increased seeking, greater short-term and long-term hyperactivity, greater short-term and long-term anxiety, greater long-term immobility, and transient but complete analgesia following fentanyl administration. We observed a very different pattern with effort escalation: hyperactivity only at low FR, minimal change in anxiety, minimal change in immobility, and increased pain for both paws and in both sham and SNI. These results at a minimum imply that brain circuits, most likely engaging mesolimbic pathways, undergo distinct adaptations with dose escalation and in effort escalation, the details of which remain to be studied.

Human studies of chronic pain patients managed with long-term opioid consumption generally demonstrate affective abnormalities such as elevated anxiety [35-38]. Our recent studies of matched chronic back pain patients either with or without long-term opioid use show opioid users demonstrate increased anxiety, depression, decreased motor abilities [39, 40], a profile that better matches the dose escalation fentanyl self-administration results.

Additionally, the majority chronic pain patients using opioids do claim pain relief [39, 40], although opioid induced hyperalgesia (OIH) following long-term opioid use is exhibited by some patients [41, 42], as well as rodents [43]. Indeed, SNI animals undergoing dose escalation experienced complete analgesia compared to the hyperalgesia seen in effort escalation. This is contrary to other findings that show analgesia in SNI mice following opioid self-administration [44]. It is possible that the time course of fentanyl in effort escalation animals was not as long-lasting as dose escalation, due to different time courses dependent on dosage. [12].Moreover, as dose escalation is the common clinical practice for managing chronic pain with opioids our mouse fentanyl dose escalation SA seems to better reflect this[15]. Yet, intriguingly we seem to have separated OIH from opioid induced analgesia based on subtle changes in opioid exposure, which suggest engagement of distinct brain circuits that remain to be uncovered.

The effects of vapor fentanyl self-administration on seeking behavior showed increased opioid motivation in females undergoing effort escalation. This is in line with research in naïve rats showing females acquire fentanyl self-administration more rapidly than males [45]. It is possible females are more sensitive to the rewarding effects of opioids, which may in turn lead to the higher opioid abuse rates in women [46]. Similar differences may also exist within the dose escalation regimen but may not have emerged given the limited sample size.

We observed an interaction of chronic pain and sex on opioid motivation, in which SNI females showed greater motivation for opioids with increasing effort. Other work with a complete Freund’s adjuvant injury led to increased intravenous fentanyl self-administration in male, but not female rats [47], while exposure to morphine shows no difference during the acquisition phase between SNI and sham animals [44]. However, authors looking at various intravenous opioids, including fentanyl, found that male rats with a spinal nerve ligation self-administered less than sham animals at lower doses, but similar amount of infusions at higher doses [12]. It is possible that the dose of opioid during effort escalation was insufficient to cause larger differences. Additionally, the effects of vapor fentanyl has not been established as much as intravenous methods, which may have different pain and sex-dependent effects on brain reward and addiction processing circuits.

During abstinence, female mice in the effort escalation regimen showed higher active port preference than male mice. The latter seems to match the human condition since in addition to higher drug use, women also show higher craving and withdrawal during abstinence [38, 48]. Another paper showed that morphine seeking behavior in extinction is only present in SNI mice, which may underline the importance of looking at cue-induced seeking in addition to acquisition [44]. There were only minor differences between SNI and sham females and males, respectively in effort escalation during cue-induced seeking. Future studies must be done to assess the interaction of sex and chronic pain on context/cue-induced seeking in dose escalation.

Overall, many factors such as session length, opioid type, pain model, and route of administration, all contribute to create lasting changes in opioid seeking behavior. Dose escalation seems advantageous to effort escalation in reflecting the human chronic pain patients managed with opioids, with or without signs of opioid use dependence. Mechanisms that would differentiate dose escalation in comparison to effort escalation potentially can provide insights into the human clinical condition.

## Notes

### Competing Interest Statement

The authors have declared no competing interest.

